# Apathy is Associated with Reduced Precision of Prior Beliefs about Action Outcomes

**DOI:** 10.1101/672113

**Authors:** Frank H. Hezemans, Noham Wolpe, James B. Rowe

**Affiliations:** MRC Cognition and Brain Sciences Unit, University of Cambridge; Department of Clinical Neurosciences, University of Cambridge

**Keywords:** apathy, motivation, goal-directed action, Bayesian, sensorimotor prediction

## Abstract

Apathy is a debilitating syndrome that is associated with reduced goal-directed behaviour. Although apathy is common and detrimental to prognosis in many neuropsychiatric diseases, its underlying mechanisms remain controversial. We propose a new model of apathy, in the context of Bayesian theories of brain function, whereby actions require predictions of their outcomes to be held with sufficient precision for ‘explaining away’ differences in sensory inputs. In this active inference model, apathy would result from reduced precision of prior beliefs about action outcomes. Healthy adults (*N*=47) performed a visuomotor task that independently manipulated physical effort and reward, and served to estimate the precision of priors. Participants’ perception of their performance was biased towards the target, which was accounted for by precise prior beliefs about action outcomes. Crucially, prior precision was negatively associated with apathy. The results support a Bayesian account of apathy, that could inform future studies of clinical populations.

## INTRODUCTION

Apathy is common, debilitating and detrimental to the prognosis in many neurological and psychiatric diseases (Lanctôt et al., 2017; Lansdall et al., 2019), but it also occurs to varying degrees in the healthy population (Ang, Lockwood, Apps, Muhammed, & Husain, 2017). Apathy is a complex construct, often decomposed into emotional, cognitive, and behavioural domains (Levy & Dubois, 2005). However, its underlying mechanisms are controversial and several accounts have been put forward for the reduction in ‘goal-directedness’ of behaviour that characterises apathy.

Behavioural economics and reinforcement learning models cast apathy primarily as a pathology of value-based decisions (Husain & Roiser, 2018). On this basis, apathetic individuals behave in ways that fail to maximise their utility, given information about the likely costs and benefits of different actions. Specifically, they exert less effort for reward (Chong, Bonnelle, & Husain, 2016), which has been attributed to deficits in dopamine-dependent reward sensitivity (Adam et al., 2013; Le Bouc et al., 2016; Muhammed et al., 2016). However, defining apathy as a lack of dopamine-dependent motivation has limitations. Current paradigms constrain action to be the consequence of a stimulus (such as a reward cue) and subsequently evaluate the action against an external reward function. This does not directly address the subject’s desire to actively fulfil their internal goals and beliefs or expectations (Gottlieb & Oudeyer, 2018). Goal-directed behaviour can alternatively be regarded as anticipatory rather than reflexive, such that actions are driven by their intended consequences (Hommel, Müsseler, Aschersleben, & Prinz, 2001).

Here we propose that apathy is directly related to the dependence of motivated behaviour on the precision of the representations of internal goals and beliefs about action outcomes. We build on the concept that brain function is a form of hierarchical Bayesian inference (Clark, 2013; Hohwy, 2013). On this basis, the brain maintains a generative model that optimises predictions of sensory inputs and minimises prediction error or ‘surprise’ (Friston, 2010; Friston, Daunizeau, Kilner, & Kiebel, 2010). Prediction error can be minimised in two ways: *passively*, by changing predictions to better fit the sensory inputs (perceptual inference), or *actively*, by performing actions to change the sensory input itself (active inference; Adams, Shipp, & Friston, 2013; Friston et al., 2010). We propose that apathetic behaviour is a disorder of active inference, within a generalised Bayesian framework.

The key question with regard to apathy is how the balance between perception and action is regulated. Under active inference theory, the precision (inverse uncertainty) of predictions and sensory input determine their relative contribution to behaviour. When predictions are held with high precision, they will be maintained even in the face of conflicting sensory input, and induce action so that the predicted and current state of the world are no longer in conflict. This means that action requires sensory attenuation: the transient down-weighting of sensory prediction errors so that expectations and goals can be fulfilled through action (Brown, Adams, Parees, Edwards, & Friston, 2013; Wolpe et al., 2016, 2018). Thus, a driving force of action is the regulation of the precision of predictions. A corollary is that low prior precision leads to a more passive behavioural state, where prediction errors are resolved by changing prior beliefs about the environment instead of by action (Friston et al., 2010, 2014).

The precision of prior beliefs can be inferred through computational modelling of behavioural data (e.g. Wolpe, Wolpert, & Rowe, 2014). This can be used to test the mechanism of individual differences in apathy, in healthy adults and clinical populations, and in relation to clinical outcomes and neural data (Adams, Huys, & Roiser, 2015). In the context of visuomotor tasks, we previously found that the precision of priors was associated with trait optimism, such that more optimistic individuals tended to have more precise priors, leading to a perceptual distortion towards better performance (Wolpe et al., 2014). Evidence from Parkinson’s disease suggests that the precision of priors is related to dopamine (Wolpe, Nombela, & Rowe, 2015).

To bring these separate lines of evidence into a common analytical framework, we hypothesised that individuals with greater apathy have less precise prior beliefs about their action outcomes. We tested this hypothesis using a visuomotor task that independently manipulated effort and reward, and from which the precision of action priors could be estimated psychophysically. We predicted that participants’ estimates of performance would be biased, in line with the integration of sensory evidence with prior beliefs about action outcomes. Using Bayesian modelling of the participants’ performance and their reported perception of performance, we estimated the precision of participants’ priors and its fluctuation across levels of effort and reward. We tested whether individual differences in apathy are related to variation in the precision of priors, and how the precision of priors depends on effort and reward.

## METHOD

### Participants

We aimed to be sufficiently powered to detect moderate associations between task metrics and apathy as follows: to detect a true correlation of *ρ* = 0.4 with *α* = .05 (two-tailed) and power of 80%, the required sample size is 46. We recruited 53 healthy adult participants to account for at least 10% data exclusions from aberrant performance profiles or technical issues. The participants had no history of a neurological or psychiatric disorder and had normal or corrected-to-normal vision. The study was approved by the Cambridge Psychology Research Ethics Committee, and all participants provided written informed consent. Participants received a standard compensation of £6 per hour and a bonus of up to £5 based on performance. Participants completed the Apathy Motivation Index (AMI; Ang et al., 2017), a questionnaire measure of apathy that is designed for the healthy adult population.

We excluded five participants whose average task performance was ≥ 3 times the median absolute deviation from the group median performance, and one participant who could not perform the force calibration appropriately. The reported analyses are therefore based on 47 participants (24 females, age range of 18–35 years, mean = 24.75, *SD* = 4.79; further demographics are given in *Table S1*).

### Task and procedure

The visuomotor task (see Figure 1) was designed to infer the precision of prior beliefs and its influence on the perception of action outcomes, under different levels of effort and reward. Participants pressed a force sensor to control the subsequent ballistic trajectory of a ‘ball’ cursor on the screen (32 pixel radius). The aim of each trial was to ‘land’ the cursor on the target (38 pixel radius). The target was either displayed close (512 pixels from left margin) or far (896 pixels from left margin) from the ball’s start position (128 pixels from left margin), such that the distance to travel corresponded to 35% (Low Effort condition) or 65% (High Effort) of each participant’s maximum force. Performance was either rewarded (in ‘points’, to be converted to cash reward after the study), or not rewarded (Reward or No Reward condition).

**Figure 1:**
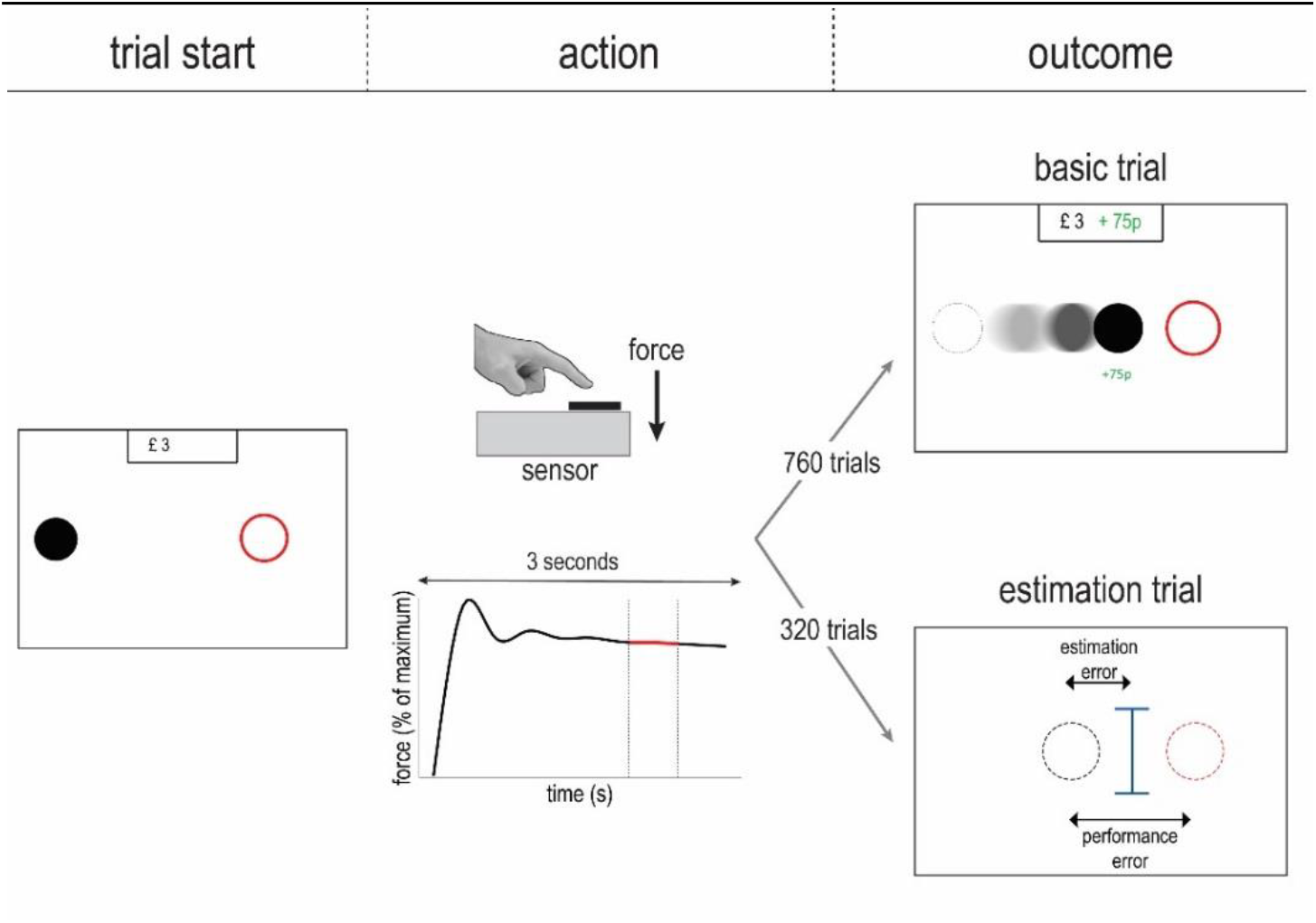
Overview of the visuomotor task. Participants performed a sustained finger press to trigger a ballistic ball trajectory, aiming it at a target. The target was either displayed close to or far from the ball’s start position, corresponding to 35% (Low Effort) or 65% (High Effort) of the participant’s maximum force. Further, participants were either given a performance-dependent monetary reward or no reward. For a minority of trials, the ball’s movement trajectory was not displayed, and participants estimated the ball’s final position with a cursor.

For each trial, participants performed a sustained finger press for 3 seconds, after which the black ball turned green to indicate that the finger could be released. Within the 3-second recording, we took the mean force from 2 to 2.5 seconds as the response (Wolpe et al., 2016). The force response determined the initial velocity of the ball. The deceleration of the ball was constant, and therefore the initial velocity (i.e. force response) uniquely determined the ball’s final position. The difference between the force response and the force needed to land the ball perfectly on target constitutes the *force error*, expressed as a percentage of the participant’s maximum force.

The task consisted of two types of trials: basic trials and estimation trials. For basic trials, participants viewed an animation of the ball’s trajectory from the start position to the final position in the direction of the target -that is, the outcome of their action. The difference between the ball’s final position and the target constitutes the *performance error*, expressed in pixels. For estimation trials, the ball’s trajectory was hidden and participants used a mouse cursor to provide their estimate of where the ball would have finished. The difference between the estimated final ball position and the true final ball position constitutes the *estimation error*, expressed in pixels. Note that the target was not displayed during the estimation procedure, and participants did not receive any feedback regarding the true final ball position. Furthermore, participants were not pre-cued about what type of trial they were engaging in. For estimation trials the ball’s animation started as usual, but after traveling 10% of the screen width, the screen turned blank and the cursor was drawn to the screen.

The experiment started with two practice blocks of 50 basic trials each, with the target in the centre of the screen (704 pixels from the left margin). In the second practice block, participants were asked to estimate their performance after viewing the full trajectory of the ball, to introduce the estimation procedure. The test phase consisted of 40 blocks of 27 trials each. We used a 2×2 full-factorial design (Low and High Effort and No Reward and Reward). In the reward condition, the maximum score of £1 was given when the ball landed perfectly on the target, and this score decreased linearly as performance error increased. To avoid confounding the effort and reward manipulations, the minimal performance required for a reward was more stringent in the Low Effort condition than in the High Effort condition (15% of the screen width from the close target versus 30% of the screen width from the far target).

There were 10 blocks of trials for each combination of effort and reward, and the blocks were ordered pseudorandomly for each participant at the start of the experiment. Each block consisted of 19 basic trials and 8 estimation trials. The trial order within each block was determined pseudorandomly, with the constraints that the first 3 trials were always basic trials, and that there could never be two consecutive estimation trials. Overall, excluding practice, participants completed 1080 trials, of which 320 were estimation trials. To reduce fatigue effects, we gave participants the opportunity to take a short break after completing a block.

#### Maximum force calibration

At the start of the task, we established each participant’s maximum force in order to normalise the effort levels between participants. This procedure consisted of 3 trials of 10 seconds each. Participants pressed with the maximum level of force they could sustain for the duration of the trial, using the index finger of their dominant hand. At the end of each trial, a sliding window function was used to select the 5 second window with the lowest force variance, and the mean force within that window was taken as the maximum force for that trial. The highest value across trials was taken as the participant’s true maximum force.

The maximum force was used to convert the force response to the ball’s initial velocity. The applied force was divided by 25% of the maximum force and then multiplied by 30% of the screen width per second. That is, pressing at 25% of one’s maximum force caused the ball to initially move at 30% of the screen width per second. To make the task less difficult under higher levels of force, we also scaled the relationship between force and initial velocity by multiplying the applied force by 0.5.

### Data analysis

#### Task performance

We preprocessed the data as follows: (i) we removed the first trial from each block to exclude any effects of switching between experimental conditions, which reduced the total number of trials from 1080 to 1040 (260 per condition); (ii) for each participant and each condition, we removed trials with a force error that was more than 3 times the median absolute deviation away from that condition’s median. On average, we removed 4 trials per condition for each participant.

We first examined the effects of effort and reward on behavioural performance. Accuracy (median force error) and variability (interquartile range, IQR, of force error) served as dependent variables in repeated measures ANOVA, with effort and reward as within-subjects factors. We report generalised eta-squared 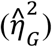 as the estimate of effect size, and we performed post-hoc Tukey’s tests to compare levels of effort and reward. We also performed Bayes factor analyses with the default ‘JZS’ prior to quantify the relative evidence in favour of a model, given the data.

#### Precision of prior beliefs

To infer the precision of prior beliefs in the perception of action outcomes, we examined participants’ estimates of their own performance as follows. For each estimation trial, we assumed that the prior and sensory evidence are Gaussian, such that the optimal estimate of the ball’s final position can be derived from Bayes’ rule (Wolpe et al., 2014):

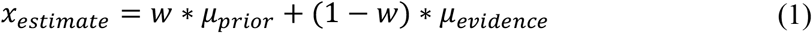

where the weighting *w* is given by:

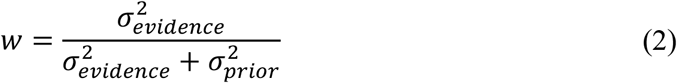

For a given estimation trial, we consider the ball’s true final ball position as the mean of the sensory evidence distribution, and the target position as the mean of the prior distribution. If a participant has no clear prior expectation regarding their performance (i.e. a ‘flat’ prior with very large variance 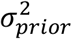), the estimate of the ball’s final position would be similar to the ball’s true final position, affected only by sensory noise with variance 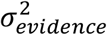. Conversely, if a participant has an exaggerated expectation of success (i.e. a prior with very small variance 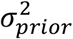), this prior would ‘overwhelm’ the sensory evidence, leading to estimates of performance that are biased towards the target relative to the true final ball position.

#### Relative weighting of priors

The first equation can be rewritten as:

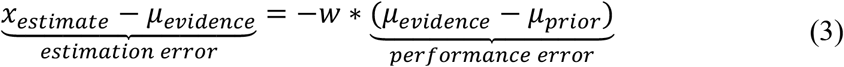

where the slope of a linear regression of estimation error by performance error characterises the weighting term *w* (Wolpe et al., 2014). A slope of −1 corresponds to full reliance on priors relative to sensory evidence, whereas a slope of 0 corresponds to a disregard of priors relative to sensory evidence.

We used linear mixed models to fit the linear relationship of estimation error by performance error. As a baseline model, we allowed the intercept and slope to vary by participants. Given that we expected performance error to depend on effort and reward, we also fit a set of models with an additional random effect term to allow for adjustments by effort and reward within each participant. Specifically, within each participant we allowed either the intercept, the slope, or both to vary by either effort, reward, or both, resulting in nine additional linear mixed models. For each model we retrieved the conditional Akaike Information Criterion (cAIC) as an approximation to the log model evidence. We selected the model with the lowest cAIC value as the most parsimonious model.

#### Modelling of prior variance

To estimate the prior variance for each participant, we fit the data with a set of hierarchical Bayesian models. The first model assumed that the prior distribution was centered on the target with unknown variance 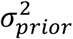, and the sensory evidence distribution was centered on the true final ball position with unknown variance 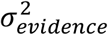. The observed estimates of the ball’s final position were then modeled as a precision-weighted combination of the prior distribution and the sensory evidence distribution (see equation 1). However, performance of the task may have been affected by computational imperfections (Stengård & van den Berg, 2019), such as perceptual shifts as a result of the ball’s rightward motion or a general bias towards the centre of visual space. The second model therefore featured an additional free parameter, ***s***, to account for directional shifts in the mean of sensory evidence:

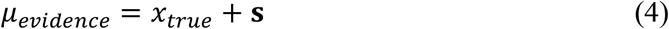

Although we consider the target as the mean of the prior distribution, participants could instead use ‘observational’ priors that reflect their actual performance distribution. We therefore additionally fit the data with a model that was similar to the first model, except the prior distribution was determined by the mean and standard deviation of each participant’s true performance on basic trials (i.e. when the true final ball position was shown).

We estimated the free parameters hierarchically: (i) parameters for individual participants were considered samples from group-level Gaussian distributions; (ii) within each participant parameters were permitted to vary between experimental conditions. Further details about the model specification are provided in Figure 5A and *Figure S1*).

We used Markov Chain Monte Carlo sampling to approximate the posterior distributions of parameters simultaneously at the level of the group, participant, and conditions. For each model we used 8 independent chains with 2000 samples, discarding the first 1000 samples as the ‘burn-in’ period. We assessed model convergence by the chains’ time series plots, and confirmed that the potential scale reduction statistic 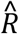 was less than 1.01 for all parameters. To identify the best model, we computed the Widely Applicable Information Criterion (WAIC) as well as approximate leave-one-out (LOO) cross-validation using Pareto smoothed importance sampling (Vehtari, Gelman, & Gabry, 2017). These estimate the pointwise predictive accuracy of a model (penalised for the effective number of parameters), using the log-likelihood evaluated at the posterior simulations of the parameter values. Our primary interest was in the participant-level estimates of prior variance, as well as the change in prior variance across levels of effort and reward.

#### Prior precision and trait apathy

We tested the relationship between model estimates of prior variance and individual differences in trait apathy. We measured trait apathy with the Apathy Motivation Index (AMI), a questionnaire measure of apathy that is suitable for the healthy population and has strong psychometric properties (Ang et al., 2017). The AMI provides a mean total score as well as mean scores for three different domains of apathy: behavioural activation, emotional sensitivity and social motivation. Each subscale consists of 6 Likert-scale items that are scored from 0 to 4, where higher scores indicate greater apathy.

As action priors correlate with performance variability and in order to rule out the effect of performance, we computed the partial correlation between trait apathy and prior precision using Pearson’s correlation, adjusted for individual differences in performance variability. Specifically, we used each participant’s standard deviation of performance error for all basic trials as an index of performance variability. We performed the partial correlation analysis separately for each outcome of the AMI. We also report the Bayes Factor for partial correlations to quantify the evidence in favour of the alternative hypothesis, given the data (Wetzels & Wagenmakers, 2012).

#### Software and equipment

The task was programmed in MATLAB (R2014a) using the Psychophysics Toolbox extensions (version 3), and were displayed on a 17-inch LCD screen (1280 x 1024 pixels). The force sensor had a sampling rate of 60 Hz and a measurement accuracy of ± 9.8 mN. Statistical analyses were implemented in R (version 3.5; R Core Team, 2018; see *Table S5* for an overview of additional packages used). The hierarchical Bayesian modeling was implemented in Stan (Carpenter et al., 2017) using the `rstan` interface package. The Method and Results sections of this paper were generated from R code using the literate programming tool `knitr`. All code, data and materials are freely available through the Open Science Framework (<link to be inserted upon acceptance for publication>).

## RESULTS

### Task performance

For each subject and each condition, we obtained a distribution of force errors (Figure 2). We examined accuracy (median force error) and variability (IQR) as a function of effort and reward (Figure 3; *Tables S2 and S3*). Accuracy was lower in the High Effort condition than in the low effort condition, as participants tended to ‘undershoot’ the target in the high effort condition (*F*_(1,46)_ = 193.46, *p* < .001, 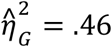). Participants were more accurate in the Reward condition than in the No Reward condition *F*_(1,46)_, *p* < .001, 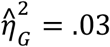) This reward effect was more pronounced in the high effort condition (effort × reward interaction: *F*_(1,46)_ = 13.32, *p* ˂ .001,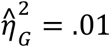). However, the Bayes Factor for the full model including an interaction effect (*BF*_10_ = 5.46 × 10^37^) was comparable to the Bayes Factor for the main effects model (*BF*_10_ = 4.22 × 10^37^), suggesting that the data do not provide clear evidence for an interaction effect (*BF*_ratio_ = 0.30).

**Figure 2:**
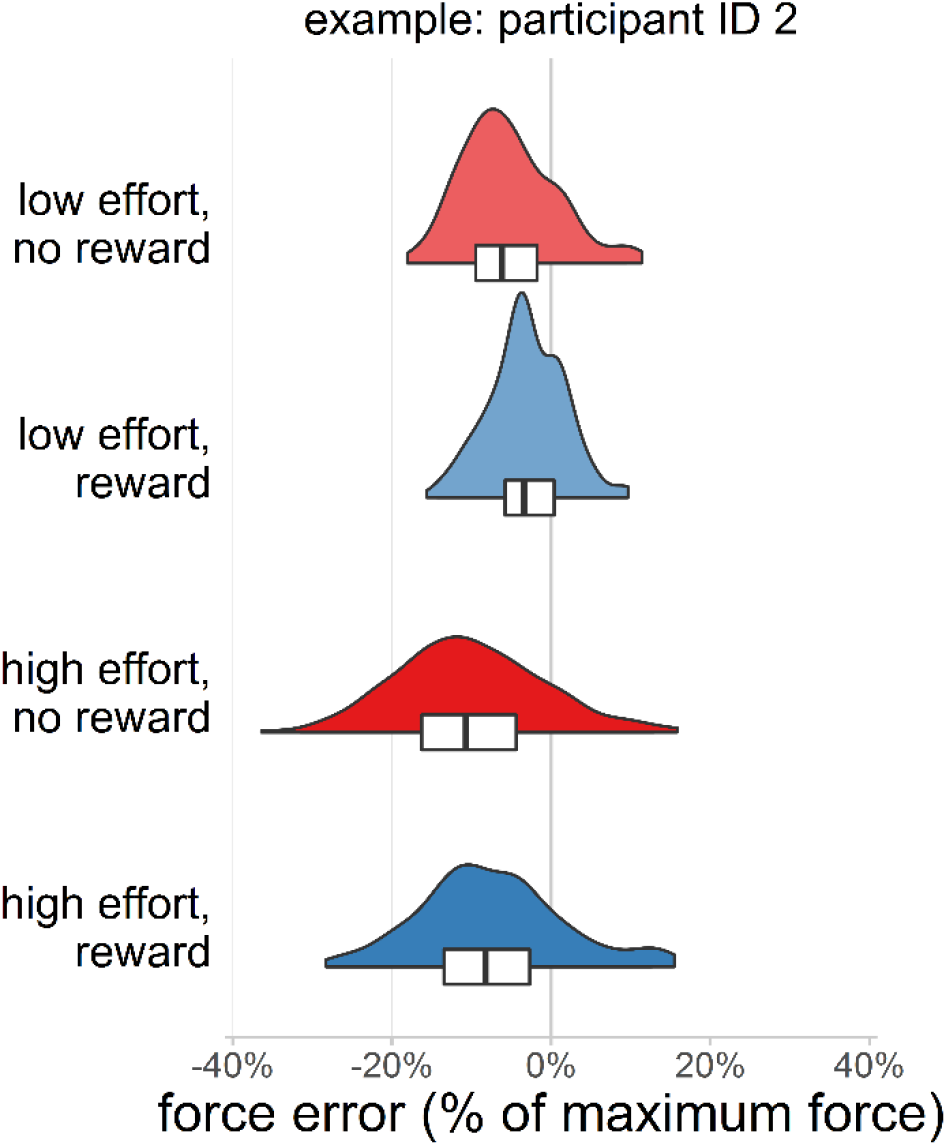
Distributions of force error by experimental conditions for a typical participant. For each distribution, the white box represents the interquartile range and the black line inside the box represents the median.

**Figure 3:**
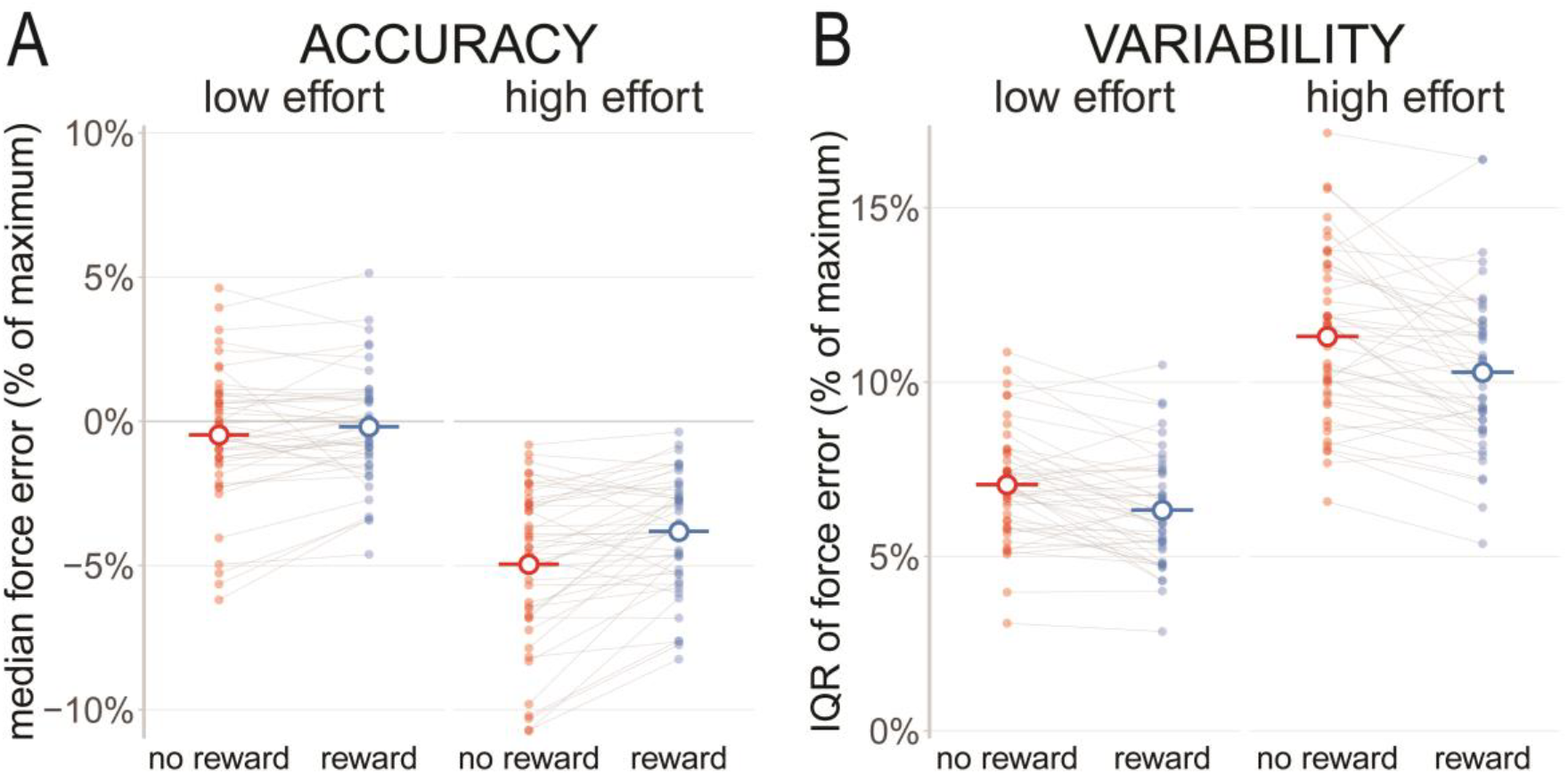
The effects of effort and reward on task accuracy (panel A) and variability (panel B). Solid dots represent the median force error (panel A) or the interquartile range of force error (panel B) for a given participant and experimental condition. The hollow dots and horizontal line segments represent the group-level mean for a given experimental condition.

Variability was greater in the High Effort condition than in the Low Effort condition(*F*_(1,46)_ = 396.35, *p* < .001,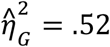). There was reduced variability in the Reward condition compared to the No Reward condition (*F*_(1,46)_ = 31.67, *p* < .001,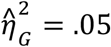), but this reward effect was not different between effort conditions (no effort × reward interaction: *F*_(1,46)_ = 1.12, *p* = .295). Bayes Factor analysis confirmed that the main effects only model (*BF*_10_ = 8.19 × 10^48^) was more likely than the full model (*BF*_10_ = 2.42 × 10^48^), providing positive evidence against an interaction effect (*BF*_ratio_ = 0.30).

Together, these results confirm that on more effortful trials, performance accuracy decreased and variability increased. In contrast, reward improved accuracy and variability in performance.

### Perception of performance

We tested whether there was a bias in the perception of action outcomes, as found in previous studies (Wolpe et al., 2015, 2014). To this end, we measured the extent to which estimates of action outcomes were biased relative to the veridical action outcomes. We first determined whether the linear relationship between estimation error and performance error varied as a function of effort, reward, or both. We fitted a set of linear mixed models that adjusted each participant’s regression intercept and slope. The ‘full model’ allowed for different intercepts and slopes by both effort and reward, and was the most likely model with an AIC difference to the next best model of 26.32 (*Table S4*). We therefore report the parameters derived from the full model.

Estimation errors tended to be biased, consistent with a prior centered on the target position (Figure 4; cf. Wolpe et al., 2014). The extent of this bias depended on performance error, as revealed by a strongly negative slope between estimation error and performance error (group-level *β* = −0.71, *CI*: [−0.75, −0.67]). Individual differences in the slope ranged from −0.9 to −0.36, confirming that all participants exhibited this estimation bias.

**Figure 4:**
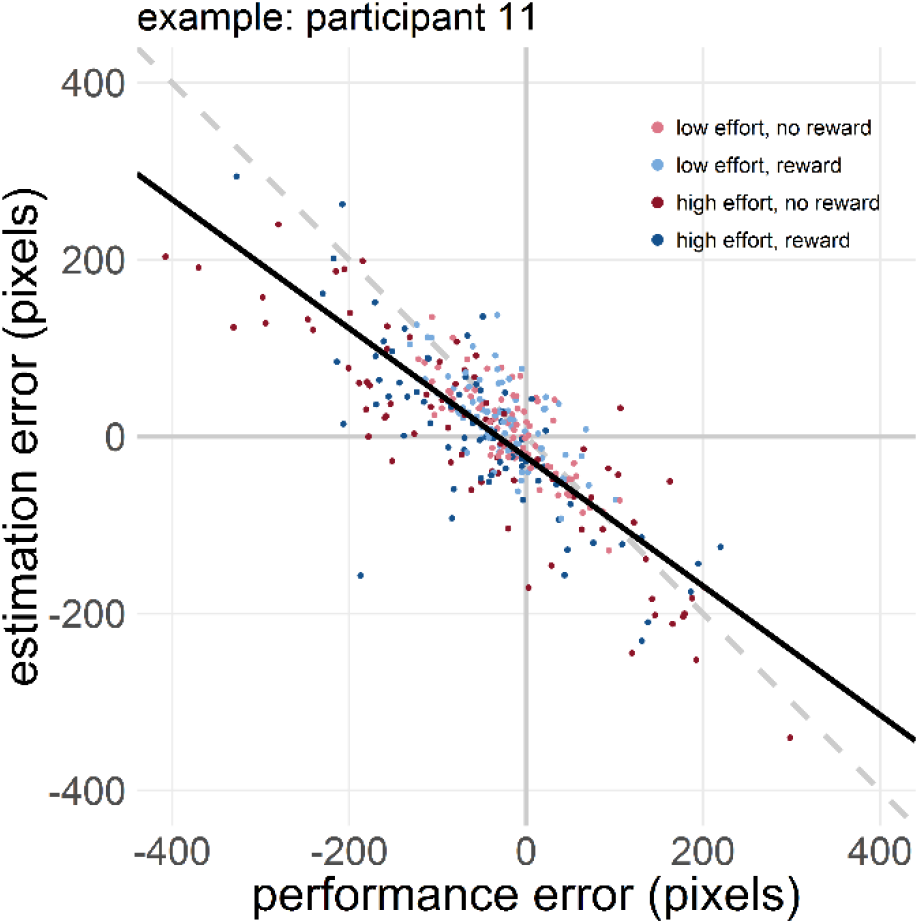
Estimation errors (difference between estimated and true ball position) plotted against performance errors (difference between true ball position and target) for a typical participant. The slope of the regression across conditions (black line) indicates the degree to which estimates of performance were biased.

### Prior precision

To test the hypothesis that trait apathy is associated with the precision (inverse of variance) of priors for the perception of outcomes, we used hierarchical Bayesian models (Figure 5A) to estimate each participant’s variance of priors. The model that best accounted for the data assumed that the prior was centered on the target position, and included a spatial shift in sensory evidence. This model was strongly preferred over a model without a sensory evidence shift (*ΔLOO*_*IC*_ = 5642.15, 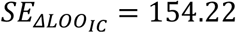 *ΔWAIC* = 5642.25, *SE*_*ΔWAIC*_ = 154.20) as well as a model with the prior determined by the mean and standard deviation of participants’ true performance (*ΔLOO*_*IC*_ = 7491.53, 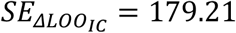 *ΔWAIC* = 7496.62, *SE*_*ΔWAIC*_ = 178.76). As illustrated in Figure 5B, there was a good agreement between the selected model’s posterior predictions and the observed data. We therefore proceeded with examining the posterior estimates of the participant-level prior standard deviation, SD.

**Figure 5:**
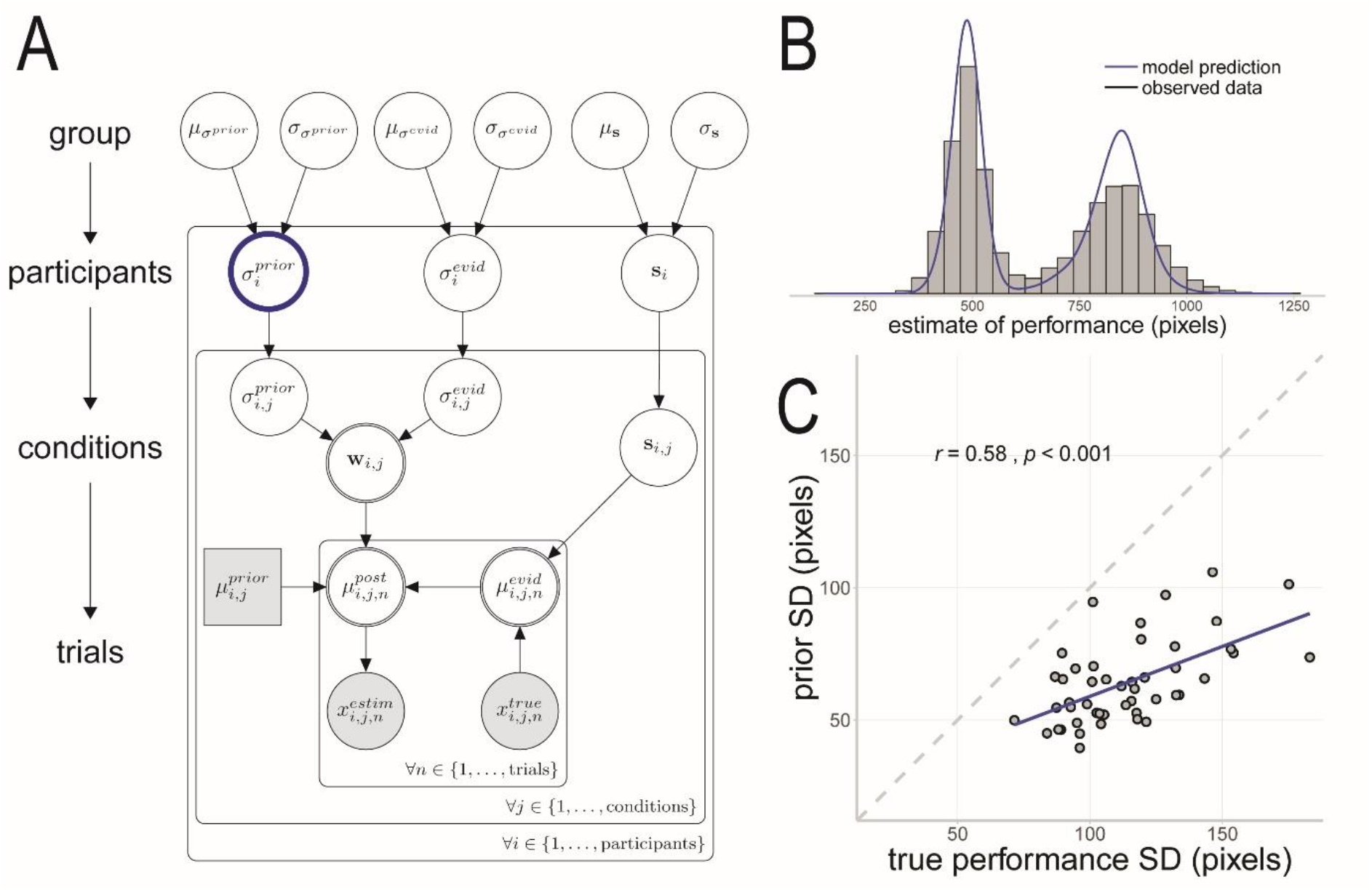
A) The best fitting Bayesian model. Shaded nodes represent observed data whereas the white nodes represent latent variables. The rectangular node represents the target position, which is a discrete variable, whereas the remaining circular nodes represent continuous variables. The double-bordered nodes represent deterministic variables that are a function of other variables without stochastic contribution. The variable of primary interest, the standard deviation of the participant-level prior, is highlighted with a blue border. B) Posterior predictive check for the best fitting model. The grey histogram represents the observed estimates of performance, and the blue density trace represents the model’s posterior predictions of estimates of performance. C) Scatterplot of the standard deviation of participant-level prior against standard deviation of performance error.

The estimates of prior SD were consistently smaller than the corresponding performance error SD (*t*_(46)_ = 17.24, *p* < .001; Figure 5C). There was also a strong correlation between prior SD and performance error SD (*r*_(45)_ = 0.58, *p* < .001). These results suggest that participants held overly precise priors that did not simply reflect the statistics of their true performance in the task.

Within participants, we allowed the prior SD to vary between experimental conditions (Figure 5A). We therefore examined prior SD as a function of effort and reward (*Figure S2*). Given that prior SD was scaled to performance SD (Figure 5C), and performance SD was strongly affected by effort and reward (Figure 3B), we normalised the prior SD to performance SD (as the ratio between prior SD and the sum of prior SD and performance SD; Wolpe et al., 2015). This normalised prior SD was smaller in the High Effort condition than in the Low Effort Condition (*F*_(1,46)_ = 66.24, *p* < .001, 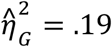). In contrast, normalised prior SD was larger in the Reward condition than in the No Reward condition (*F*_(1,46)_ = 7.16, *p* = .010,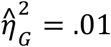). There was no significant interaction effect between effort and reward (*F*_(1,46)_ = 1.25, *p* = .269). Bayes Factor analysis confirmed that the main effects only model (*BF*_10_ = 3.72 × 10^15^) was more likely than the full model (*BF*_10_ = 1.21 × 10^15^), providing positive evidence against an interaction effect (*BF*_*ratio*_= 0.32).

To test whether the variance of prior beliefs about action outcomes is associated with trait apathy, we used partial correlations of the participant-level estimates of prior SD and the Apathy Motivation Index scores, adjusting for task performance variability. There was a significant correlation between prior SD and the AMI behavioural activation subscale (*r*_(45)_ = 0.36, *p* = .012, Holm-Bonferroni corrected *p* = .050; *BF*_10_ = 3.67; Figure 6). The association was positive, suggesting that individuals who were more apathetic had reduced prior precision. There were no significant partial correlations with the AMI total score (*r*_(45)_ = 0.18, *p* = .220; *BF*_10_ = 0.37), emotional sensitivity subscale (*r*_(45)_ = 0.12, *p* = .413; *BF*_10_ = 0.24), or social motivation subscale (*r*_(45)_ = −0.11, *p* = .458; *BF*_10_ = 0.23).

**Figure 6:**
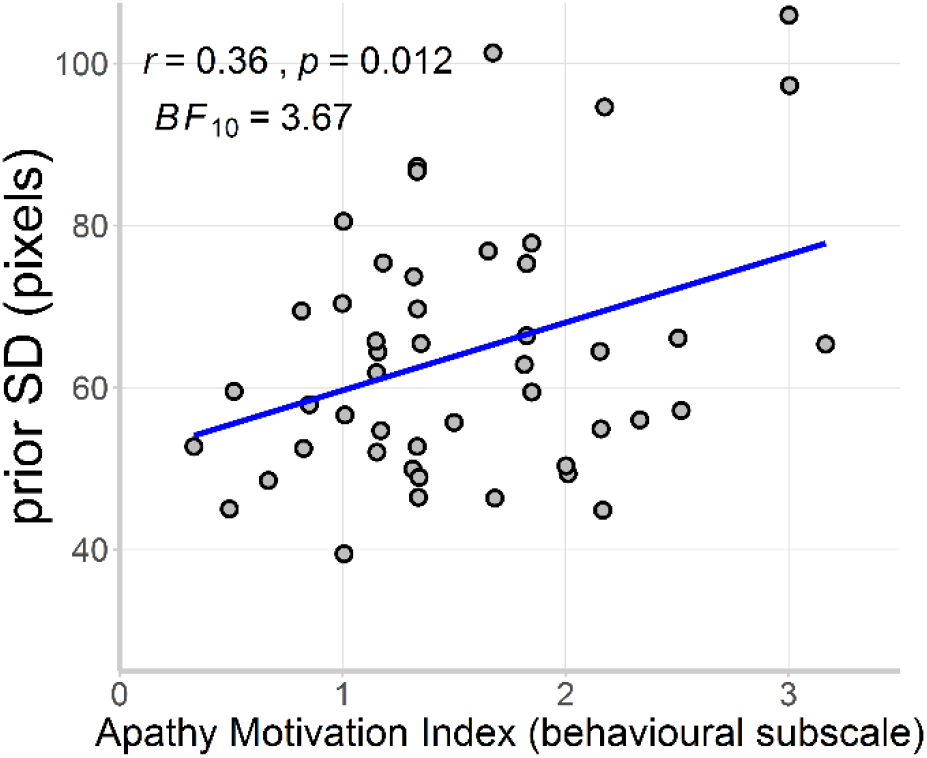
Scatterplot of the standard deviation of participant-level prior against the Apathy Motivation Index behavioural activation subscale. For illustration purposes, identical questionnaire scores are jittered.

We explored whether changes in prior SD between experimental conditions were associated with trait apathy, but found no evidence for such relationships. The difference between reward conditions in normalised prior SD was not significantly associated with the AMI total score (*β* = −0.07, *p* = .639), behavioural activation subscale (*β* = 0.19, *p* = .168), emotional sensitivity subscale (*β* = −0.13, *p* = .358), or social motivation subscale (*β* = −0.22, *p* = .120). These associations also did not depend on effort, as there were no significant interaction effects between effort and the change in normalised prior SD by reward (all *p* values ≥ .194). Bayes Factor analysis confirmed that the intercept only model was more likely than the full model for all AMI scales, providing positive evidence against associations with trait apathy (all *BF*_10_ ≤ 0.04). In terms of effort, the difference between the Low Effort and High Effort conditions in normalised prior SD was not associated with the AMI total score (*β* = −0.01, *p* = .965), behavioural activation subscale (*β* = 0.15, *p* = .305), emotional sensitivity subscale (*β* = −0.08, *p* = .590), or social motivation subscale (*β* = −0.10, *p* = .498), and there were no significant interaction effects between reward and the change in normalised prior SD by effort (all *p* values ≥ .306). Bayes Factor analysis confirmed that the intercept only model was more likely than the full model for all AMI scales, providing positive evidence against associations with trait apathy (all *BF*_10_ ≤ 0.02).

## DISCUSSION

The principal result of this study is that higher trait apathy is associated with lower precision of prior beliefs about action outcomes. In the context of effortful, goal-directed actions, we confirmed that people’s perception of performance was biased, relative to the veridical action outcomes. Participants’ estimation of their action outcomes were explained by ‘overly precise’ priors that do not simply reflect the statistics of performance. The variability of these priors was associated with trait apathy, such that more apathetic individuals tended to have less precise priors of action outcomes.

These results are consistent with a Bayesian framework of brain function, in which the brain engages in active inference on the causes of sensory inputs. Central to this model is that prior beliefs and sensory input are combined in a *precision-weighted* manner, so that more precise (i.e. less uncertain) information plays a stronger role in shaping action and perception. A hypothesis emerging from this framework is that a loss of prior precision leads to bradykinesia and a loss of goal-directed behaviour (e.g. Friston et al., 2010, 2014). Our results support this hypothesis.

There is evidence that apathetic individuals are less incentivised by rewards, particularly when those rewards require investment of effort. Although reduced motivation for reward certainly can contribute to apathy (Adam et al., 2013), this mechanism does not fully explain the multifaceted nature of apathy in patient groups (Lansdall et al., 2017). For example, even in the absence of external prompts (such as a reward cue), apathetic patients often have difficulty in self-generating motor patterns, over and above blunted affect or cognitive dysfunction (Levy & Dubois, 2005). Such ‘auto-activation’ symptoms have previously been formalised as a failure to reach a necessary activation threshold for a response (Zhang et al., 2016). Patients were surprisingly biased in favour of performing an action, but were subsequently impaired at translating this prior preference into an observed response, as indicated by a strongly reduced rate of accumulation to threshold (Zhang et al., 2016). We suggest that the diversity of symptoms associated with apathy can be understood as different expressions of a common underlying pathology: a reduction in the precision of prior beliefs about action outcomes.

Although estimates of prior precision within participants changed significantly between levels of effort and reward, the amount of change between conditions did not depend on trait apathy. This suggests that participants’ overall prior for action reflects their higher-level beliefs and motivations related to trait apathy, whereas trail-to-trial changes to prior precision in light of task demands reflect lower-level mechanisms of sensorimotor prediction. In the current experiment, reward decreased the precision of priors relative to the true performance distribution. Such a strategy would promote learning in the context of expected reward (since the posterior belief will be weighted more towards the evidence than the prior), in contrast to the motivational advantage of the illusion of superiority that occurs in non-rewarded trials (when the posterior belief of success is weighted more towards the prior than the evidence). In other words, the reward manipulation facilitates learning from sensory evidence, so that trial-to-trial performance errors can be used to improve task performance.

Neuropathologies associated with apathy provide insights into the functional anatomy and candidate mechanisms of the abnormal precision of priors. Clinical apathy is associated with disruptions to frontal-subcortical circuits that are involved in self-initiated, goal-directed behaviour (Levy & Dubois, 2005). Lesions of the prefrontal cortices have long been known to impair goal-directed behaviour (Luria, 1995), and apathy is an important feature of neurodegenerative diseases affecting prefrontal regions (Passamonti, Lansdall, & Rowe, 2018). Lesion and neuroimaging studies have also implicated the anterior cingulate cortex and basal ganglia in apathy (Le Heron, Apps, & Husain, 2017; Levy & Dubois, 2005). Thus, current evidence suggests that apathy follows a disruption to fronto-striatal brain circuits.

Changes in these fronto-striatal circuits have been implicated in controlling the relative precision of predictions and sensory input (Dayan & Yu, 2006; Friston et al., 2014; Moran et al., 2013). In Parkinson’s disease, the severe depletion of striatal dopamine is associated with a loss of sensory attenuation and presence of apathy, both consequences of impaired active inference (Drui et al., 2014; Macerollo et al., 2016; Santangelo et al., 2015; Wolpe et al., 2018). Indeed, individual differences in the degree of sensory attenuation were negatively related to disease severity, but positively related to dopamine medication dose (Wolpe et al., 2018). We hypothesise that neuromodulatory deficits in patients can cause a loss of prior precision relative to the sensory input, which subsequently leads to apathy. Furthermore, higher order prior beliefs about desired outcomes may fail to appropriately contextualise lower-level representations about sensory input due to structural and functional abnormalities in prefrontal and temporal brain regions (Rittman et al., 2019). Further work is required to establish the synaptic and molecular basis of aberrant precision, aided by the parameterisation of the precision of an individual’s prior.

There are limitations to this study. First, we examined individual differences in trait apathy in the healthy population, and the generalisation to apathetic clinical disorders remains to be proven. A dimensional approach assumes that the mechanisms underlying normal variation are the same mechanisms which underlie clinical disorders (Cuthbert & Insel, 2013) but we recognise that apathetic patients might be qualitatively different to controls. Second, our primary results are correlational and therefore do not directly demonstrate causal mechanisms. Our results do not in themselves prove whether the precision of priors is a cause or consequence of trait apathy. Future work can adopt our approach to study the induction of apathy in the context of neurosurgical or temporary focal brain lesions (e.g. through transcranial magnetic stimulation) or pharmacological manipulations (e.g. Adam et al., 2013; Le Bouc et al., 2016). Third, we cannot comment on the variations in functional anatomy or connectivity that may determine the precision of priors in our cohort. Finally, the Bayesian framework for computational models enables relative evidences to be compared formally (Adams et al., 2015), but only between members of the subset of models tested.

In conclusion, our study suggests that apathy is associated with poor precision of prior beliefs about action outcomes. We propose that apathy can be understood as a failure to assign the necessary precision to prior beliefs about one’s action outcomes, necessary for self-initiated movement, leading to an apparent ‘acceptance’ of the state of the world. This can be understood as satisfying an intended goal in the absence of the actual action necessary to achieve it. This approach paves the way to a common framework for understanding the causes of apathy in neurological and psychiatric disorders, and a target for novel treatment strategies.

## AUTHOR CONTRIBUTIONS

All authors contributed to the design of the study and task paradigm. F. H. Hezemans programmed the task, and collected and analysed the data under the supervision of N. Wolpe and J. B. Rowe. All authors contributed to the interpretation of the findings. F. H. Hezemans drafted the manuscript, and N. Wolpe and J. B. Rowe provided critical revisions. All authors approved the final version of the manuscript for submission.

## ACKNOWLEDGMENTS

We thank David Hayes for building the force sensor.

## OPEN PRACTICES STATEMENT

The experiment reported in this article was not formally preregistered, but the sample size was based on an *a priori* power analysis. The de-identified raw data, MATLAB experiment code and R analysis code are freely available through the Open Science Framework and can be found at <link to be inserted upon acceptance for publication>.

## SUPPLEMENTAL MATERIAL

**Figure S1:**
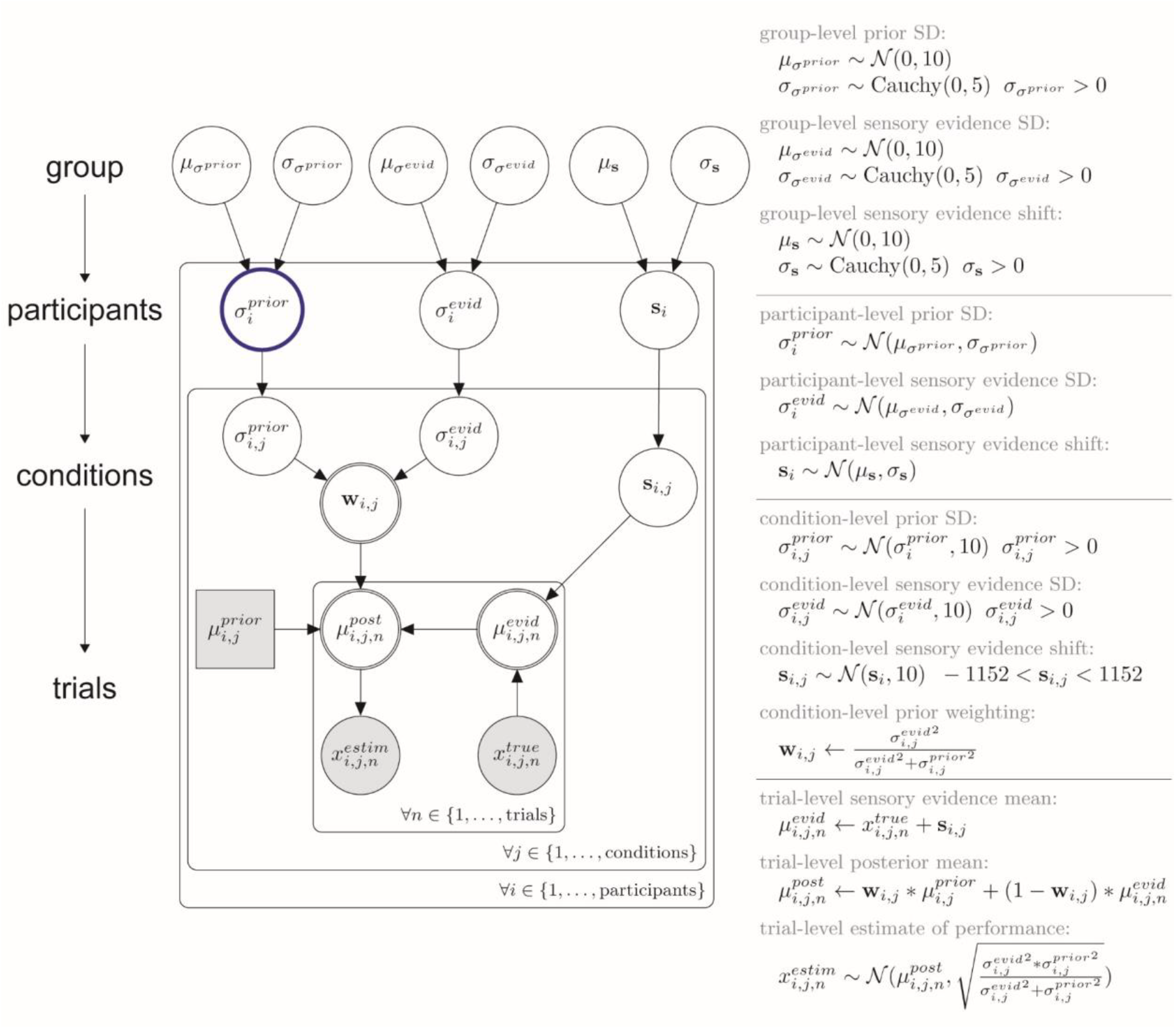
The best fitting Bayesian model (as in Figure 5A), including the parameters’ sampling statements and functional dependencies. Participant-level parameters were sampled from latent group-level parameters, and within participants parameters were permitted to vary between conditions. Condition-level parameters were then used to compute a precision-weighted combination of the prior and trial-level sensory evidence. This trial-level posterior mean served as the trial-level predicted estimate of performance. The model is represented in plate notation: shaded nodes represent observed data whereas white nodes represent latent variables; rectangular nodes represent discrete variables whereas circular nodes represent continuous variables; and double-bordered white nodes represent deterministic variables whereas single-bordered white nodes represent stochastic variables.

**Figure S2:**
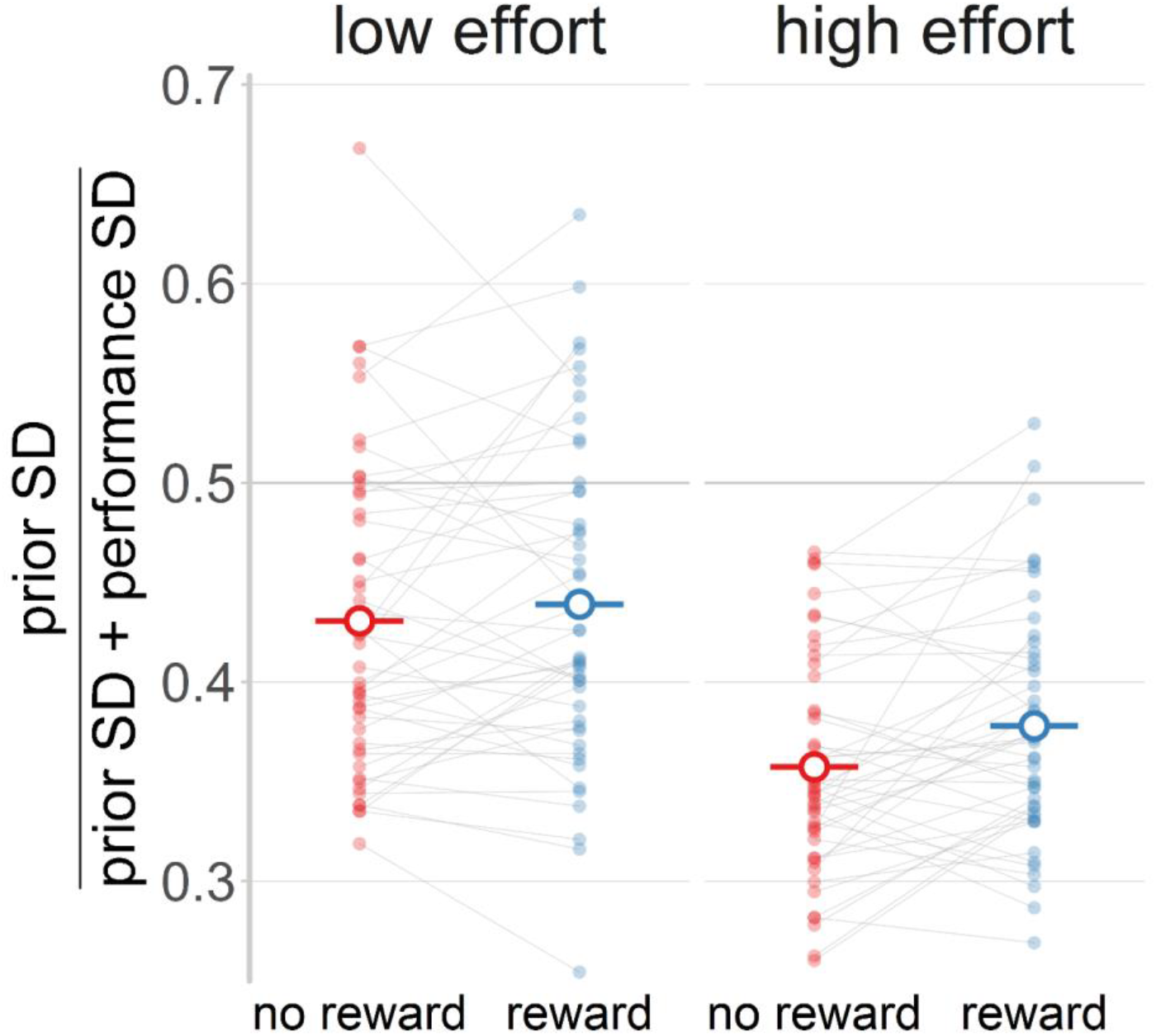
The effects of effort and reward on the standard deviation of the prior, normalised to the standard deviation of performance error (Wolpe, Nombela, & Rowe, 2015). Values smaller than 0.5 represent priors that are more precise than the corresponding performance distribution. Solid dots represent individual participants, whereas the hollow dots and horizontal line segments represent the group-level mean for a given experimental condition.

**Table S1:** Descriptive statistics of sample demographics and trait apathy

|  | **mean** | **SD** | **range** |
| --- | --- | --- | --- |
| age | 24.23 | 4.84 | 18-35 |
| education |  |  |  |
| years<sup>a</sup> | 17.30 | 3.12 | 13-26 |
| degree<sup>b</sup> | 1.94 | 0.89 | 1-4 |
| Apathy Motivation Index<sup>c</sup> |  |  |  |
| total | 1.34 | 0.42 | 0.5-2.39 |
| behavioural activation | 1.54 | 0.66 | 0.33-3.17 |
| social motivation | 1.48 | 0.65 | 0.33-2.83 |
| emotional sensitivity | 1.00 | 0.54 | 0.17-2.17 |

*Note:* 47 participants (24 females).
<sup>a</sup> total years of formal education, including everything after kindergarten.
<sup>b</sup> highest obtained degree categorised according to the British education system: 0 = GCSE (General Certificate of Secondary Education), 1 = A Levels (General Certificate of Secondary Education Advanced Level), 2 = undergraduate degree, 3 = graduate degree, 4 = postgraduate / doctorate degree.
<sup>c</sup> scored on a Likert scale from 0 to 4, with higher scores indicating greater apathy (Ang, Lockwood, Apps, Muhammed, & Husain, 2017).

**Table S2:**
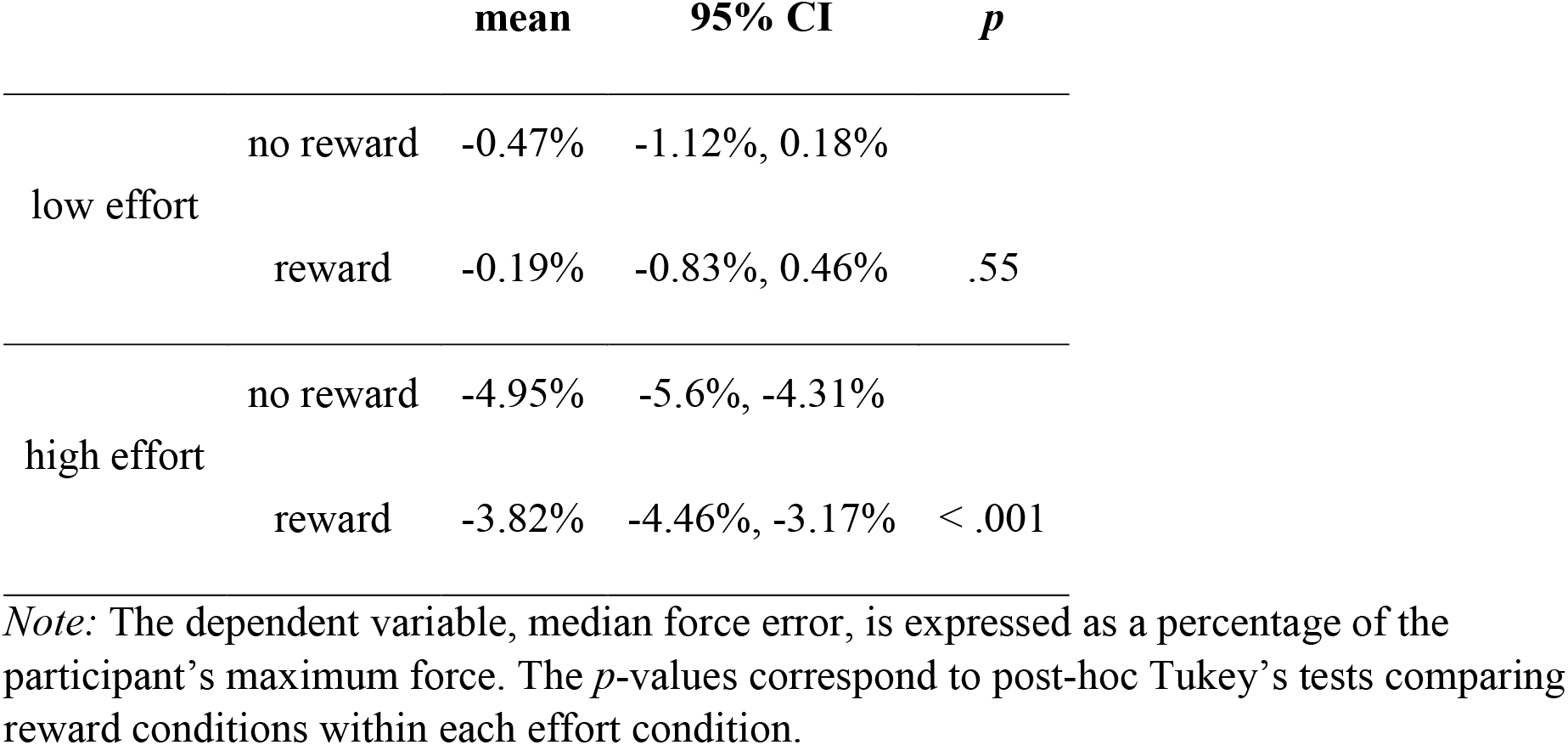
Descriptive statistics of accuracy (median force error) for each experimental condition

**Table S3:**
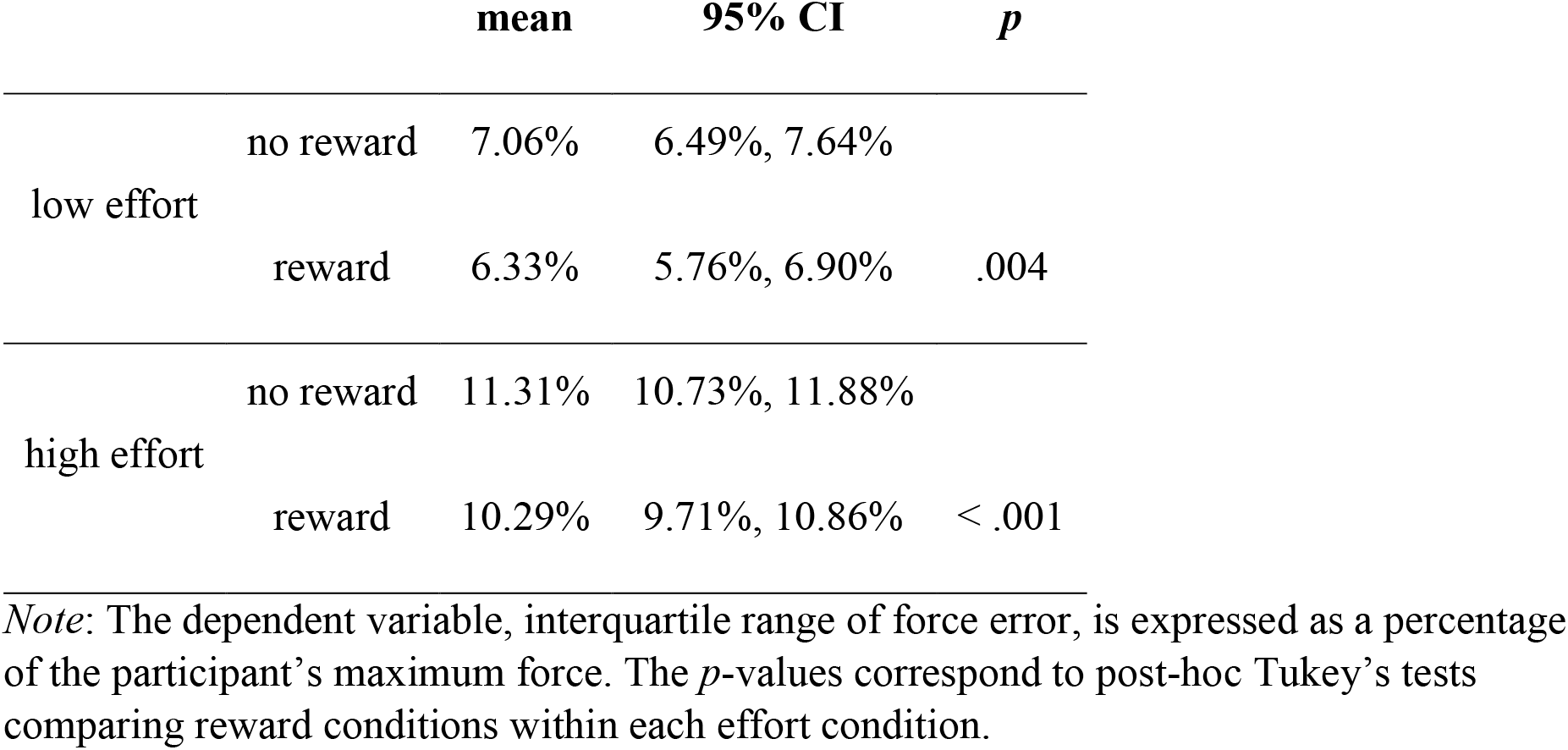
Descriptive statistics of variability (interquartile range of force error) for each experimental condition

**Table S4:** Log model evidence for linear mixed models of estimation error by performance error

| | cAIC | ΔcAIC |
| --- | --- | --- |
| basic | 30017.32 | 1873.01 |
| varying intercepts |  |  |
| effort | 28349.94 | 205.63 |
| reward | 29900.64 | 1756.34 |
| effort & reward | 28170.63 | 26.32 |
| varying slopes |  |  |
| effort | 29865.84 | 1721.53 |
| reward | 29947.07 | 1802.76 |
| effort & reward | 29817.52 | 1673.21 |
| varying intercepts & slopes |  |  |
| effort | 28342.20 | 197.89 |
| reward | 29858.90 | 1714.59 |
| **effort & reward** | **28144.31** | **0.00** |

*Note:* cAIC = conditional Akaike Information Criterion (Greven & Kneib, 2010), $\Delta$ cAIC = [cAIC – min(cAIC)]. The ‘basic’ model allowed for varying intercepts and slopes between participants, but did not take into account the experimental conditions of effort and reward. The nine remaining models allowed for further adjustments to the intercept, slope, or both by effort, reward, or both. The favoured model, with varying intercepts and slopes by effort and reward, is highlighted in bold.

**Table S5:** R packages used for statistical analysis

| **package** | **usage** | **citation** |
| --- | --- | --- |
| tidyverse (version 1.2.1) | data organisation and visualisation | (Wickham, 2017) |
| pwr (version 1.2-2) | power calculations for correlation test | (Champely, 2018) |
| afex (version 0.21-2) | ANOVA | (Singmann, Bolker, Westfall, & Aust, 2017) |
| BayesFactor (version 0.9.12-4.2) | Bayes factor for ANOVA and regression models | (Morey & Rouder, 2018) |
| emmeans (version 1.2.2) | estimated marginal means and post-hoc Tukey's tests | (Lenth, 2018) |
| lme4 (version 1.1-20) | linear mixed models | (Bates, Mächler, Bolker, & Walker, 2015) |
| cAIC4 (version 0.5) | conditional Akaike information criterion for linear mixed models | (Säfken, Rügamer, Kneib, & Greven, 2018) |
| merTools (version 0.4.1) | extract and organise parameter estimates from linear mixed model | (Knowles & Frederick, 2018) |
| rstan (version 2.18.2) | hierarchical Bayesian modelling | (Stan Development Team, 2018) |
| loo (version 2.0.0) | approximate leave-one-out cross-validation and widely applicable information criterion for Bayesian models | (Vehtari, Gabry, Yao, & Gelman, 2018) |
| tidybayes (version 1.0.4) | extract and organise parameter estimates from Bayesian model | (Kay, 2019) |
| knitr (version 1.20) | generate methods and results sections from R code | (Xie, 2018) |

